# Probing the role of macromolecular crowding in cell volume regulation using fluorescence anisotropy

**DOI:** 10.1101/2022.04.07.487477

**Authors:** Parijat Biswas, Dipanjan Ray, Subhamoy Jana, Jibitesh Das, Bidisha Sinha, Deepak Kumar Sinha

## Abstract

Cytoplasmic macromolecular crowding (MMC) influences multiple cellular functions. Using fluorescence anisotropy of EGFP, we explore the homeostasis of MMC in the cytoplasm and its role in cell volume regulation. Individual cells of different lineages maintain a distinctly unique average cytoplasmic MMC for considerably long durations. Despite diffusion, actin cytoskeleton facilitates spatially heterogeneous MMC in the cytoplasm. While hypertonic-stress triggers regulatory volume increase (RVI), other methods of increasing cytoplasmic MMC fail to start RVI, suggesting that cells lack capabilities for MMC homeostasis. During spreading or microtubule-depolymerized state, cells neglect changes in cytoplasmic MMC to undergo volume regulation. Inhibition of TNFR1 increases the membrane tension and deprives cells of their ability to undergo volume regulation. Current understanding contemplates that cytoplasmic MMC, the mechanical state of the plasma membrane or the actin cytoskeleton, could be sensors for cell volume regulation. Our observations establish the irrelevance of cytoplasmic MMC and the significance of plasma membrane tension in setting off the cell volume regulation machinery.

## Introduction

The cytoplasm and the lumen of many organelles contain different macromolecules in remarkably high densities. Cytoplasmic protein densities may range from 50-400 mg/mL, depending on the cell lineage (Fulton, 1982; Cayley *et al*., 1991). Cytoplasmic macromolecules primarily comprise proteins (56%), nucleic acids (23%), lipids (9%), polysaccharides (3%) (Kohata and Miyoshi, 2020). Consequently, the fraction of excluded volume of the macromolecules becomes notable compared to the total volume of the cytoplasm. This entropic effect is also described as the thermodynamic state of macromolecular crowding (MMC) (Minton, 1981; Ellis, 2001), which affects a host of physicochemical processes in the cytoplasm such as diffusion (Rashid *et al*., 2015), active transport (Nettesheim *et al*., 2020), protein-ligand binding kinetics (Minton, 2001; Köhn *et al*., 2021), enzyme-substrate reaction rates (Thoke, Bagatolli and Olsen, 2018; Wilcox, Chung and Slade, 2021), macromolecular self-assembly (André and Spruijt, 2020; Schreck, Bridstrup and Yuan, 2020), protein folding (Ådén and Wittung-Stafshede, 2014) and post-translational modifications (Darling and Uversky, 2018), to name a few. Consequently, it led to a hypothesis that cells may regulate their cytoplasm MMC to control biochemical processes. Despite the high relevance of cytoplasmic MMC, the literature lacks experimental study on cellular ability for homeostasis of cytoplasmic MMC (Lin and Amir, 2018). Thus, more information is needed concerning the temporal dynamics and population heterogeneity of cytoplasmic MMC in eukaryotic cells. Cytoplasmic MMC and cell volume are inversely related; therefore, MMC is hypothesized to be the thermodynamic condition that cell volume regulation machinery utilizes as a volume sensor (Minton, Colclasure and Parker, 1992; Burg, 2000; Al-Habori, 2001). However, other factors, such as the mechanical state of the plasma membrane, can alternatively act as sensors for cell volume regulation (Strange, 2004; Hoffmann, Lambert and Pedersen, 2009; Gonzalez *et al*., 2018; Adar and Safran, 2020). Some of the critical factors that alter MMC in the cytoplasm are translation, degradation of proteins, and cytoskeleton disassembly. It is unclear whether or not cells tightly maintain MMC in its cytoplasm for relatively long durations. Thus, the investigation of cytoplasmic MMC is valuable and requires a reliable tool for its dynamic characterization in living cells.

In this manuscript, we establish that steady state Fluorescence Anisotropy of EGFP (*r_EGFP_*) is a reliable indicator of cytoplasmic MMC, with a high dynamic range virtually insensitive to variations in intracellular pH, salt concentration, small-molecule concentration, and exhibit reversibility. Using *r_EGFP_*, we investigate i) spatiotemporal dynamics and ii) the ability of the cells to enforce the homeostasis of cytoplasmic MMC at the cellular level, and also explore the link between MMC and cell volume regulation.

## Results and discussion

### EGFP as a sensor for Macromolecular Crowding

We study the effect of macromolecular crowder concentrations on the fluorescence anisotropy of EGFP (*r_EGFP_*)(**Fig 1A**). The value of *r_EGFP_* rises linearly with increasing concentrations of BSA in the mM range. Polysucrose, small organic molecules, and salts induce sufficient change in *r_EGFP_* only at a very high, non-physiological concentration range (**Fig 1B**). According to the Perrin equation: *r* = *r*_0_/(1 + *τ/θ_C_*), where, *θ_C_* = *ηV/k_B_T*, the observed change in *r_EGFP_*, can be explained by the effect of MMC on fluorophore lifetime (*τ*), solution viscosity (*η*), and intrinsic anisotropy (*r*_0_). Increasing MMC increases both the solution viscosity *η* (Rashid *et al*., 2015) and the refractive index ‘*n*’, leading to a decrease in *τ*, according to the Strickler-Berg equation (1/*τ* ∝ *n*^2^) (Strickler and Berg, 1962). The rotational correlation time (*θ_C_*) of EGFP in PBS (≈14ns) (Novikov *et al*., 2017) is significantly longer than its *τ* (≈2.6ns) (**Fig S1**), hence further increase in *η* is expected to cause a negligible change in *r_EGFP_* (Suhling, Davis and Phillips, 2002; Suhling *et al*., 2002). Therefore, we conclude that the observed increase in *r_EGFP_* (**Fig 1A, Fig 1B**) in crowded solutions is caused mainly by the reduction in *τ* (**Fig 1C**) and increase in *r*_0_ (**Fig S1B**). To establish this, we compare the fluorescence anisotropy measurement of EGFP (*r_EGFP_*) with that of fluorescein (*r_FLRSCN_*) in 80-90% glycerol solutions where fluorescein’s*θ_C_* is comparable to its *τ* (Devauges *et al*., 2012). For these solutions, viscosity (*η*) increases significantly by 264%, whereas the refractive index (*n*) changes by a minuscule amount (1 %) (*CRC Handbook of Chemistry and Physics, 85th Edition - Google Books*, no date, pp. 8– 66). The shape of the *r_EGFP_* and *r_FLRSCN_* plots in **Fig 1D** establish that increase of solution *η* has a negligible effect on *r_EGFP_*. To verify the reliability of *r_EGFP_* as an indicator of MMC in the cell cytoplasm, we further explore its dependence on pH and EGFP concentration. **Fig 1E** establishes that *r_EGFP_* is independent of variations in its own concentration, and **Fig 1F** shows that *r_EGFP_* is independent of pH at the physiological range. Therefore, cell to cell variations in EGFP expression level, cytosolic pH, salt concentrations, or the small molecule crowder concentrations will not alter the measured *r_EGFP_*, making it a reliable indicator of cytoplasmic MMC.

**Fig 1:**
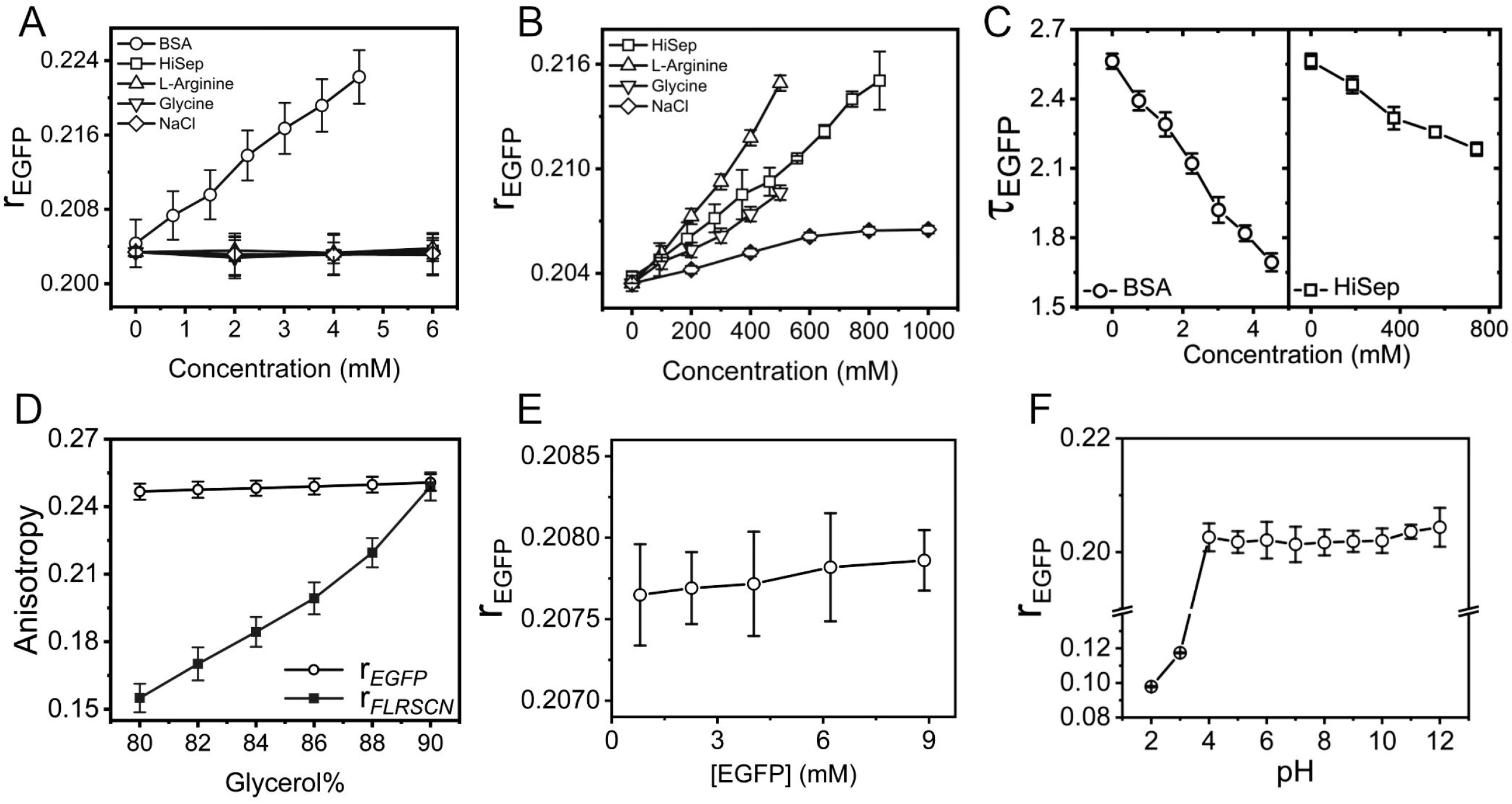
Steady state fluorescence anisotropy of EGFP as a sensor for MMC. **(A)** *r_EGFP_* increases linearly with increasing protein (BSA) and other crowder concentrations in Hepes buffer (pH 7.6). **(B)** *r_EGFP_* increases at very high, non-physiological concentrations concentration of small crowding molecules. **(C)** The fluorescence lifetime of EGFP (*τ_EGFP_*)decreases with crowder BSA and HiSep concentration. **(D)** Comparison between the *r_EGFP_* and *r_FLRSCN_* in glycerol solutions of increasing concentration. **(E)** Value of *r_EGFP_* stays relatively constant with increasing concentrations of EGFP. **(F)** *r_EGFP_* in Hepes buffers of increasing pH range (6-8).

### Cells’ ability to enforce RVI is cell lineage dependent

We directly probe the efficacy of *r_EGFP_* as a sensor for cytoplasmic MMC in cells of multiple lineages. Hypertonic shock induces rapid water efflux from cells, thus increasing the cytosolic MMC (Ho, 2006; Petelenz-Kurdziel *et al*., 2011; Miermont *et al*., 2013). On exposing EGFP expressing NIH/3T3 fibroblasts (NIH/3T3-EGFP) to dextrose supplemented hypertonic condition (950mOsm), the value of *r_EGFP_* rises, as shown in **Fig 2A**. Time-lapse measurement of *r_EGFP_* (**Fig S2A**) establishes that the cytoplasmic MMC in NIH/3T3-EGFP cells increases rapidly within 80 seconds in hypertonic conditions. Subsequently, the cells maintain the increased cytosolic MMC for at least up to 2 hours (**Fig 2B**). As expected, hypotonic shock (~170mOsm) reduces the cytoplasmic MMC (**Fig 2C, S2B**). However, cells can counteract the effect of hypotonic shock within 2 hours through regulatory volume decrease (RVD), as evident from the *r_EGFP_* and cell spread area trajectory (**Fig 2C**, **S2B**)(Roffay *et al*., 2021). The observed increase of *r_EGFP_* due to increased cytoplasmic MMC is caused primarily by the decrement of its fluorescence lifetime, *τ_EGFP_* (**Fig 2D**, **S2C**)(Suhling *et al*., 2002). While 950mOsm hypertonic shock with dextrose causes about a 20% increase in*r_BGFP_*, without recovery (**Fig 2B**), 950mOsm shock with NaCl causes a smaller change (~10%) in *r_EGFP_* (**Fig S2D**), because the rapid exchange of Cl^-^ and Na^+^ ions can alleviate the adverse effects of excessive osmotic imbalance (Vallés *et al*., 2015; Serra *et al*., 2021). Reversing the osmotic imbalance (950mOsm) with isotonic DMEM causes immediate reversal of *r_EGFP_* to pre-shock value (**Fig 2E**). Thus, the MMC sensing property of EGFP exhibits reversibility. Further exposing NIH/3T3-EGFP to varying degrees of osmotic imbalances causes a gradually larger increase in *r_EGFP_* (**Fig 2F**). These observations suggest that measurement of *r_EGFP_* permits fast, precise monitoring of a significantly large degree of changes in the cytoplasmic MMC. The rise in MMC due to hypertonic imbalance is either maintained (for ≥550mOsm) or is gradually restored to (for ≤500mOsm) within 30 minutes of hypertonic shock via regulatory volume increase (RVI) (Jalihal *et al*., 2020). Since EGFP localizes both in the cytoplasm and the nucleoplasm, we investigate the impact of enhanced cytoplasmic MMC on the mobility of cytoplasmic and nucleoplasmic proteins using FRAP. **Fig S2E** indicates that gradually higher degrees of hypertonicity, decrease the mobility of EGFP monotonically in the cytoplasm and the nucleoplasm. As expected, hypertonic shock with a non-metabolizable analog of dextrose, mannitol, has an identical impact on NIH/3T3 cells (**Fig S2F**). Our studies indicate that different cell lineages, such as RAW 264.7, HeLa, and MDA-MB-231, maintain a different level of average cytoplasmic MMC (**Fig S2H**). Investigation of RVI establishes cell-type dependent response to identical hypertonic shock (**Fig 2G-i, ii**). Interestingly, RAW 264.7 macrophages exhibit an approximate rise of 14% and 17% in *r_EGFP_* within 10 minutes exposed to 400mOsm (**Fig 2G-i**, **Fig S2G-i**) or 500mOsm (**Fig 2G-ii**, **Fig S2G-ii**) dextrose, respectively. It is a significantly larger change than observed in NIH/3T3 cells (~2% and ~6%, respectively). Further, unlike NIH/3T3 fibroblasts, RAW 264.7 and MDA-MB-231 cells fail to restore the effect of osmotic imbalance endogenously during 30 minutes time intervals. However, the response of HeLa cells to hypertonic shock is comparable to that of the NIH/3T3 fibroblasts (**Fig 2G-i, ii**). The osmotic imbalance causes changes in cytoplasmic MMC via fast efflux/influx of water. We employ other methods that perturb the cytoplasmic MMC without inducing quick water exchange- (i) by treating cells with Heclin, an inhibitor of HECT E3 ubiquitin ligase (Mund *et al*., 2014), to reduce protein degradation, thereby increasing cytosolic protein concentration, (ii) by using Cycloheximide to inhibit protein translation (Siegel and Sisler, 1963), thus reducing cytosolic protein concentration, and (iii) exposing cells to heat shock, causing widespread cytoplasmic protein degradation (Parag, Raboy and Kulka, 1987). Cells incubated with Heclin show a significant rise in cytoplasmic MMC, whereas Cycloheximide treatment causes a small but significant reduction of cytoplasmic MMC. Heat shock causes a considerable decrease in *r_EGFP_* values (**Fig 2H**). Using Bradford assay, we independently validate **Fig 2H** by estimating total protein mass per cell in response to Heclin, Cycloheximide, and heat shock (**Fig 2H, inset**). To understand whether the higher cytoplasmic MMC in RAW 264.7 cells (**Fig S2H**) disarm them of their ability to undergo RVI **(Fig 2G)**, we expose NIH/3T3-EGFP cells pretreated with Heclin to 500mOsm hypertonic shock. Increasing MMC in NIH/3T3-EGFP cells does not deprive it of its ability for RVI (**Fig S2I**), suggesting the ability of the cells to enforce RVI is independent of its cytoplasmic MMC and inherent to its cell type.

**Fig 2:**
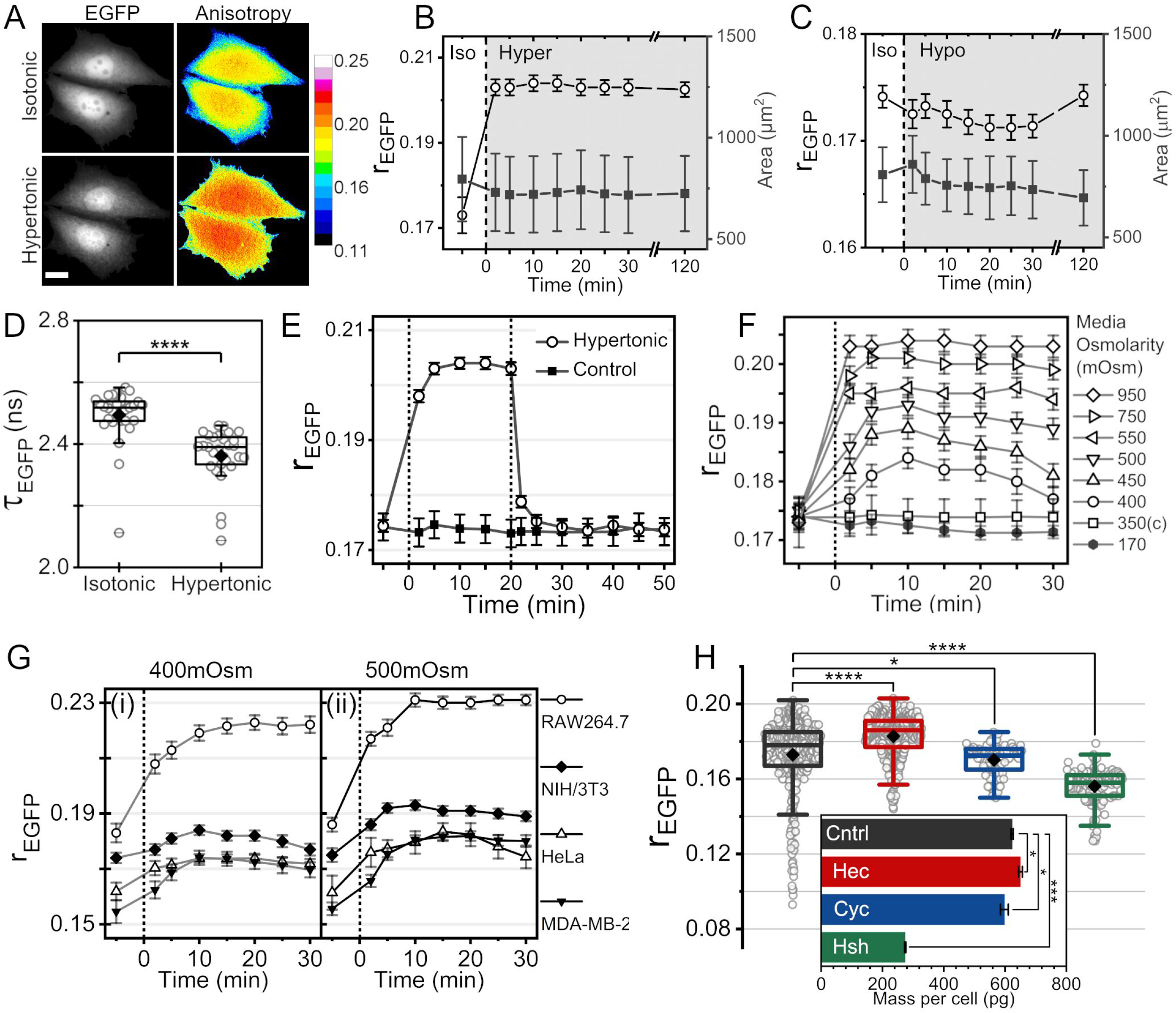
RVI of cells in hypertonic media is cell lineage dependent. **(A)** Intensity and *r_EGFP_* map of NIH/3T3-EGFP exposed to hypertonic conditions (950mOsm). The Colour bar on the right shows the corresponding anisotropy values. Scale bar=15μm. **(B)** Quantification of *r_EGFP_* and cell spread area of NIH/3T3-EGFP exposed to 950mOsm hypertonicity. The black dotted line indicates the time of changing media tonicity. The white area represents the isotonic condition; the grey area represents the hypertonic condition. **(C)** Quantification of *r_EGFP_* and cell spread area of NIH/3T3-EGFP exposed to 170mOsm hypotonicity. The black dotted line indicates the time of changing media tonicity. The white area represents the isotonic condition; the grey area represents the hypotonic condition. **(D)** Quantification of fluorescence lifetime in NIH/3T3-EGFP exposed to the hypertonic condition. **(E)** Comparison of *r_EGFP_* in NIH/3T3-EGFP exposed to hypertonic condition (950mOsm) for 20min, with isotonic control. The black dotted line at t=0min indicates the time of changing tonicity to hypertonic condition, and t=20min shows the time of reverting media tonicity to isotonic condition. **(F)** Quantification of *r_EGFP_* in NIH/3T3-EGFP cells exposed to culture media of different tonicities (170mOsm-950mOsm). The black dotted line indicates the time of changing media tonicity **(G)** Comparison of *r_EGFP_* in different cell lines upon exposure to hypertonic conditions. **(H)** Comparison of *r_EGFP_* in NIH/3T3-EGFP upon treatment with Heclin (70μM, 4h; in red), Cycloheximide (1μM, 4h; in blue) or heat shock (42°C, 1h; in green). Inset represents mean protein mass per cell under the same treatment.

### Actin filaments facilitate spatially heterogeneous MMC in the cytoplasm

The representative *r_EGFP_* images of NIH/3T3-EGFP in isotonic conditions (**Fig 3A**) establishes that the cytoplasmic MMC is non-uniform at a few microns’ length scales. In a significant fraction of different cell lineages, namely, Hela (~54%), NIH/3T3 (~23%), RAW 264.7 (~16%), and MDA-MB-231 (~14%), the level of nucleoplasmic MMC is higher than the neighbouring perinuclear cytoplasm. Additionally, the cortical regions have lower levels of cytoplasmic MMC than the perinuclear regions of the cytoplasm. Further, **Fig 3A** establishes that the MMC in the lamellipodial region is lower than the rest of the cell body, suggesting differential MMC levels within the cytoplasm. Time-lapse investigation of cells that generate new lamellipodial extensions further establishes that the MMC in newly developed lamellipodial cytoplasm is lower than the rest of the cell body (**Sup. Vid. 1**), which confirms previous rheology measurements (Laurent *et al*., 2005). In regions where the thickness of cells is comparable to the Z-resolution of the microscope, the cells show lower *r_EGFP_* values. Therefore, in addition to lifetime imaging (**Fig S2C**), we validate the differential cytoplasmic MMC in the lamellipodial and the perinuclear cytoplasm independently by alternate methods. Using FRAP, we compare the translational diffusion rate of EGFP in lamellipodial and the perinuclear cytoplasm (**Fig 3B i**). Higher MMC leads to an increase of solution viscosity (Rashid *et al*., 2015); however, the FRAP results in **Fig 3B-i** exhibit an indiscernible difference in the viscosities of the lamellipodial (η=4.2±1.1cP) and perinuclear (η=4.8±1.6cP) cytoplasm. We conclude that the changes in the bulk viscosity caused by an alteration in MMC are probe-size dependent and cannot be measured with a probe whose size is comparable to or smaller than the crowder molecules (Fabry *et al*., 2001; Wong *et al*., 2004). We verify this *in vitro* by measuring the viscosity of protein solutions (BSA: 0mM, 2.26mM, and 4.52mM) with two different methods. Estimation of solution viscosity with FRAP of EGFP (size of EGFP is comparable to BSA) gives η=2.73±1.1cP for BSA free Hepes buffer, η=3.37±1.32cP for 2.26mM BSA, and η=3.51±3.64cP for 4.52mM BSA at 20°C (**Fig S3A-i**). However, estimation of solution viscosity using single particle tracking method with a probe bigger than the crowder molecules, i.e., 200nm fluorescent microspheres (**Fig S3A-ii**), provides 1.48±2.72cP for Hepes buffer (0mM BSA), 3.09±5.41cP for BSA 2.26mM, and 26.02±4.07cP for BSA 4.56mM at 20°C, which are significantly different from the FRAP measurements, and agrees with reported bulk viscosity measurements (Yadav, Shire and Kalonia, 2011). It establishes that a 30%(w/v) increase in MMC using BSA affects the translational thermal motion (Mean Squared Displacement or MSD) of probe EGFP molecules by only 22%; as a result, only a small change in solution viscosity is observed by FRAP (**Fig S3A-i**). However, a similar increase in MMC using BSA reduces the translational thermal motion of bigger-sized probe particles by 94%, as observed by single particle tracking (**Fig S3A-ii**). The comparison of MSD of 200nm probe particles reveals that the lamellipodial cytoplasm has a lower crowding (*η* =2.22±1.63cP) in comparison to the perinuclear cytoplasm (*η* =18.56±15.82cP) (**Fig 3B-ii**). Interestingly, the sharp boundary that separates the low MMC cytoplasmic region at the cortex from the rest of the cytoplasm is often associated with actin filaments (shown by the arrows in **Fig 3C**). Thus, we hypothesize that the actin stress fibers are crucial in maintaining the sharp difference in cytoplasmic MMC (**Fig 3C**). Disruption of actin filaments in cells leads to homogenization of the cytoplasmic MMC (**Fig S3B-i**), establishing the importance of actin fibers in maintaining the boundary. Disruption of microtubules and the intermediate filament vimentin leads to an insignificant change in the spatial distribution of cytoplasmic MMC (**Fig S3B-ii, iii**). Disassembly of actin leads to a significant rise in average cytoplasmic MMC compared to disassembly of microtubules and vimentin (**Fig 3D**). It is probably because disassembly of F-actin generates a significantly larger concentration of corresponding actin monomers compared to microtubules and intermediate filaments (Liebermeister *et al*., 2014; Pegoraro, Janmey and Weitz, 2017; Loiodice *et al*., 2019). Having established the non-uniformity in cytoplasmic MMC distribution, we investigate whether hypertonic conditions cause a spatially uniform rise of cytoplasmic MMC (**Fig 3E**). Interestingly, water expulsion in response to hypertonic shock is non-uniform at the cortex, compared to the perinuclear regions. The lamellipodial edges exhibit a higher amount of water expulsion than the rest of the cell boundaries.This observation agrees with our claim that lamellipodia have lower MMC than the rest of the cytoplasm; as a result, it expels more water, leading to a larger increase in cytoplasmic MMC. Additionally, the literature suggests that lamellipodia have more aquaporins, and thus can expel more water than non-lamellipodial edges of the cells (Karlsson *et al*., 2013; Papadopoulos and Saadoun, 2015; Direito *et al*., 2016).

**Fig 3:**
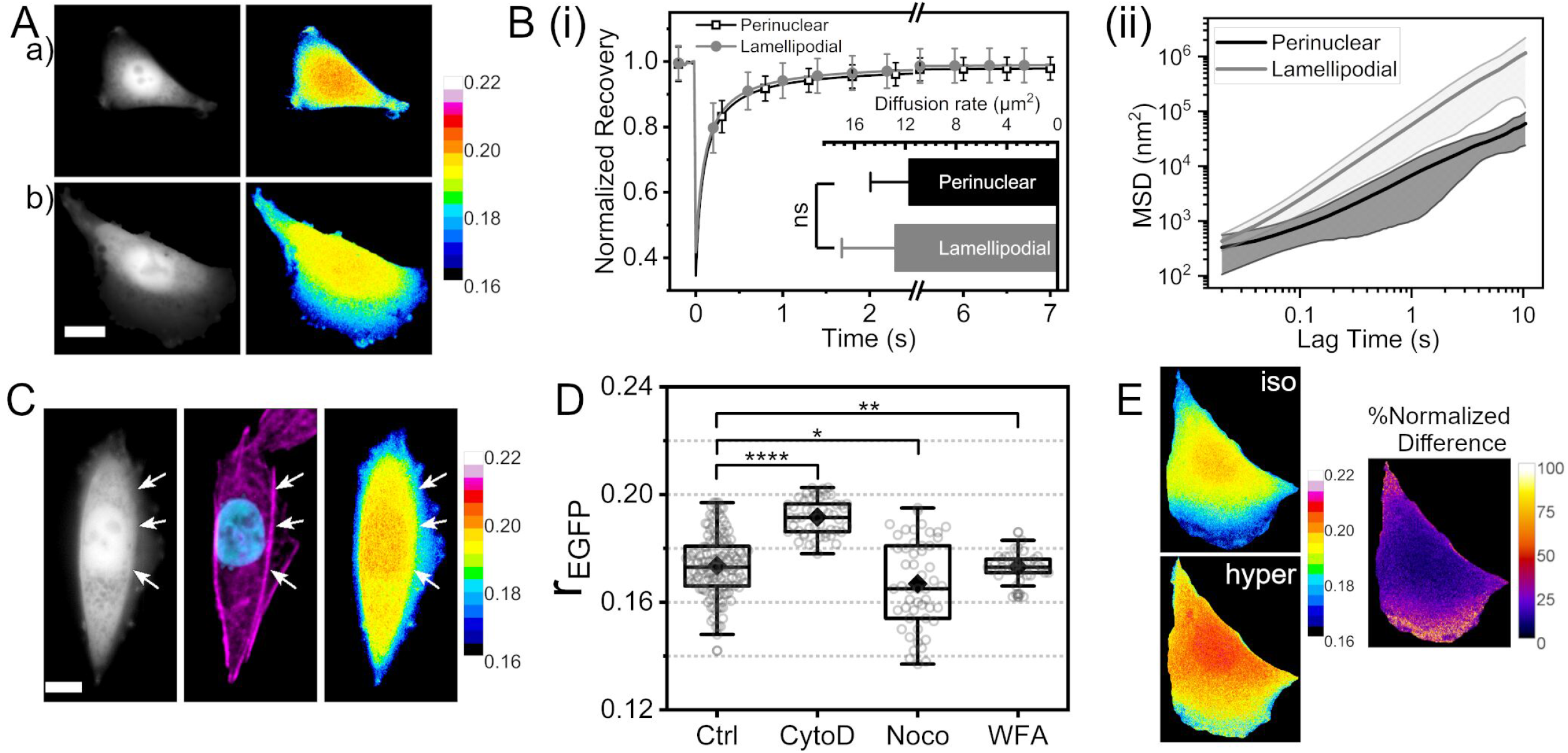
Despite diffusion, actin filaments hold spatially heterogeneous MMC in the cytoplasm. **(A)** Fluorescence Intensity image and *r_EGFP_* map of representative NIH/3T3-EGFP cells showing spatially varying cytoplasmic MMC; **a** represents a cell lacking lamellipodia and having higher nucleoplasmic MMC; **b** shows a cell with prominent lamellipodia. The Colour bar on the right shows the corresponding anisotropy values. Scale bar=15μm. **(B-i)** FRAP analysis of translational mobility of EGFP in the lamellipodial and perinuclear regions. **(B-ii)** Mean Squared Displacement (MSD) from single particle tracking of fluorescent microspheres in the lamellipodia and perinuclear regions. **(C)** Highlighting the spatial variation of MMC governed by the actin cytoskeleton. The dashed white line separates regions of lower MMC from higher MMC regulated by the actin density. First column-EGFP intensity; Middle column-phalloidin-Alexa488 stained actin (magenta) and Hoechst 33342 stained nucleus (cyan); Third column-*r_BGFP_* map. Scale bar=15μm. **(D)** Cytoplasmic *r_EGFP_* after disruption of actin by Cytochalasin D (2μM,1h), microtubules by Nocodazole (20μM,1h) and vimentin by Withaferin A (3μM,3h). **(E)** Spatially differential response of the cytoplasm to hypertonic shock (500mosm). Lamellipodial regions show a higher change in *r_EGFP_* than perinuclear regions as visible from the % normalized difference map (third column). Scale bar=15μm.

### Hypertonic stress-induced non-specific protein aggregation in the cytoplasm is independent of the increase in MMC

The macromolecular concentration in the cytoplasm should be inversely proportional to its volume if the number of macromolecules remains unchanged. Thus, we investigate the dependence of cytoplasmic *r_EGFP_* on the cell volume in different hypertonic solutions (**Fig 4A**). However, the shape of the measured *r_EGFP_* vs cell volume (*V*) curve in **Fig 4A** is different from the expected relation: 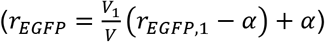. It suggests that, in addition to volume change, other cellular processes may alter *r_EGFP_* in response to the hypertonic condition. The shape of the curve in **Fig 4A** suggests that there is either an increase in the total number of cytoplasmic macromolecular crowders or a change in the effective crowder size. However, measurement of protein mass per cell (**Fig 4B**) establishes that there is no measurable increase in the amount of cytosolic protein content in response to hypertonic shock. Thus, we explore other possibilities like the reorganization of the cytoplasmic macromolecular crowders in response to hypertonic shock that changes the effective crowder size, causing a further rise of *r_EGFP_* in **Fig 4A** (indicated by the arrow). We mimic the reorganization of proteins in the cell cytoplasm by fixing NIH/3T3-EGFP with 4% para-formaldehyde. Fixation causes covalent links between neighbouring protein molecules, increasing the effective crowder size around EGFP molecules. **Fig 4C** compares the cytoplasmic *r_EGFP_* of cells before and after fixation, establishing that increase in the effective crowder size is an additional factor that can increase *r_EGFP_* along with the change in cell volume. Super-resolution imaging of cytoplasmic EGFP in cells exposed to different hypertonic strengths confirms protein aggregation in the cytoplasm (**Fig 4D**). However, other methods of increasing cytoplasmic MMC (Heclin treatment, **Fig 2H**) do not lead to noticeable protein aggregation (**Fig 4E**). Literature also suggests that multivalent proteins form larger aggregates in response to increased MMC via self-condensation (Shin and Brangwynne, 2017; Watanabe *et al*., 2018; Jalihal *et al*., 2020, 2021; Keber *et al*., 2021). While free EGFP depicts a non-specific weaker aggregation in the cytoplasm, NFκβ, a protein with DNA binding domain (Yan *et al*., 2012; Smith *et al*., 2019; Vonderach *et al*., 2019), shows globular self aggregation in response to hypertonic shock in NIH/3T3 cells expressing p65-EGFP, a subunit of the NFκβ complex (**Fig S4A-i**). However, in Heclin treated NIH/3T3, NFκβ shows drastically massive, irregularly shaped aggregates (**Fig S4A-ii**) compared to the globular condensates from hypertonic shock. According to previous reports (Irianto *et al*., 2013), hypertonic conditions cause DNA condensation in the nucleus (**Fig S4B-i**). However, Heclin mediated increase of cytoplasmic MMC has a non-measurable impact on chromatin condensation (**Fig S4B-ii**). Therefore, we conclude that the cytoplasmic MMC is not responsible for the condensation of chromatin.

**Fig 4:**
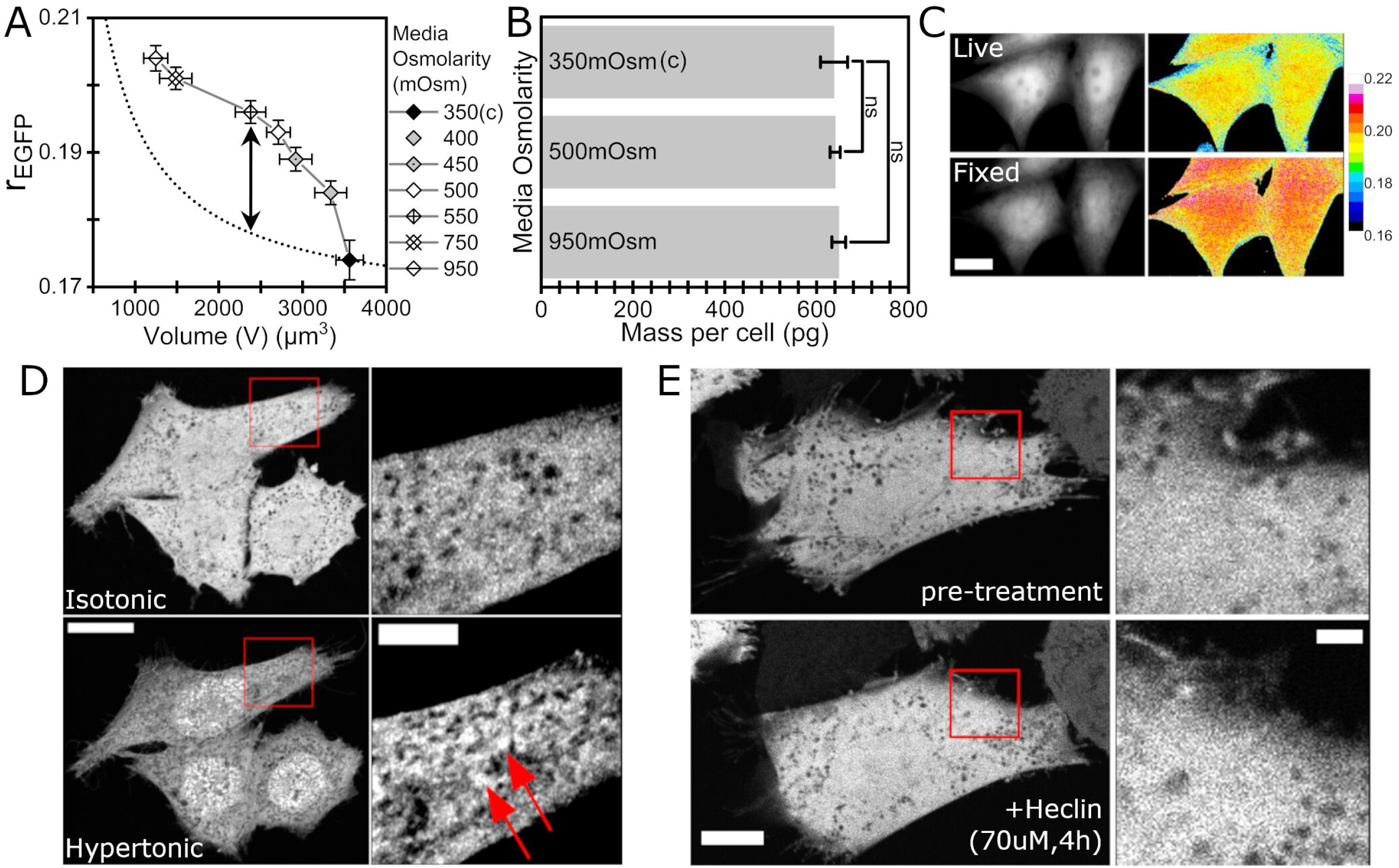
Hypertonic stress induces rapid and widespread submicron protein aggregation. **(A)** Quantification of *r_EGFP_* and cell volume at different hypertonic conditions. **(B)** Estimation of total protein mass per cell at different hypertonic conditions. **(C)** Comparison of *r_EGFP_* in live and fixed cells. Scale bar=15μm. **(D)** AiryScan super-resolution imaging of NIH/3T3-EGFP reveals submicron clustering of EGFP under hypertonic conditions (950mOsm). Scale bar: main=15μm, inset=5μm. **(E)** AiryScan super-resolution imaging of NIH/3T3-EGFP before and after Heclin treatment shows no apparent aggregation of EGFP. Scale bar: main=15μm, inset=2μm.

### Cell spreading activates RVI

We observe a weak negative correlation (*r*=-0.386) between the spread area and the corresponding cytoplasmic MMC (*r_EGFP_*) in NIH/3T3-EGFP (**Fig 5A**). It suggests a positive correlation between the volume of the cytoplasm (*V*) and the cells’ spread area (*A*). The spread area of NIH/3T3 exhibits a positive, linear correlation (r=0.909) to the cell volume; *V* = 4.2*A* + 86.8 (**Fig 5B-i**). This correlation could arise either from the cell cycle or from the extent of cell spreading independent of its cell cycle stage. To resolve this, we explore the dynamics of cytoplasmic MMC during cell spreading in NIH/3T3-EGFP.

**Fig 5:**
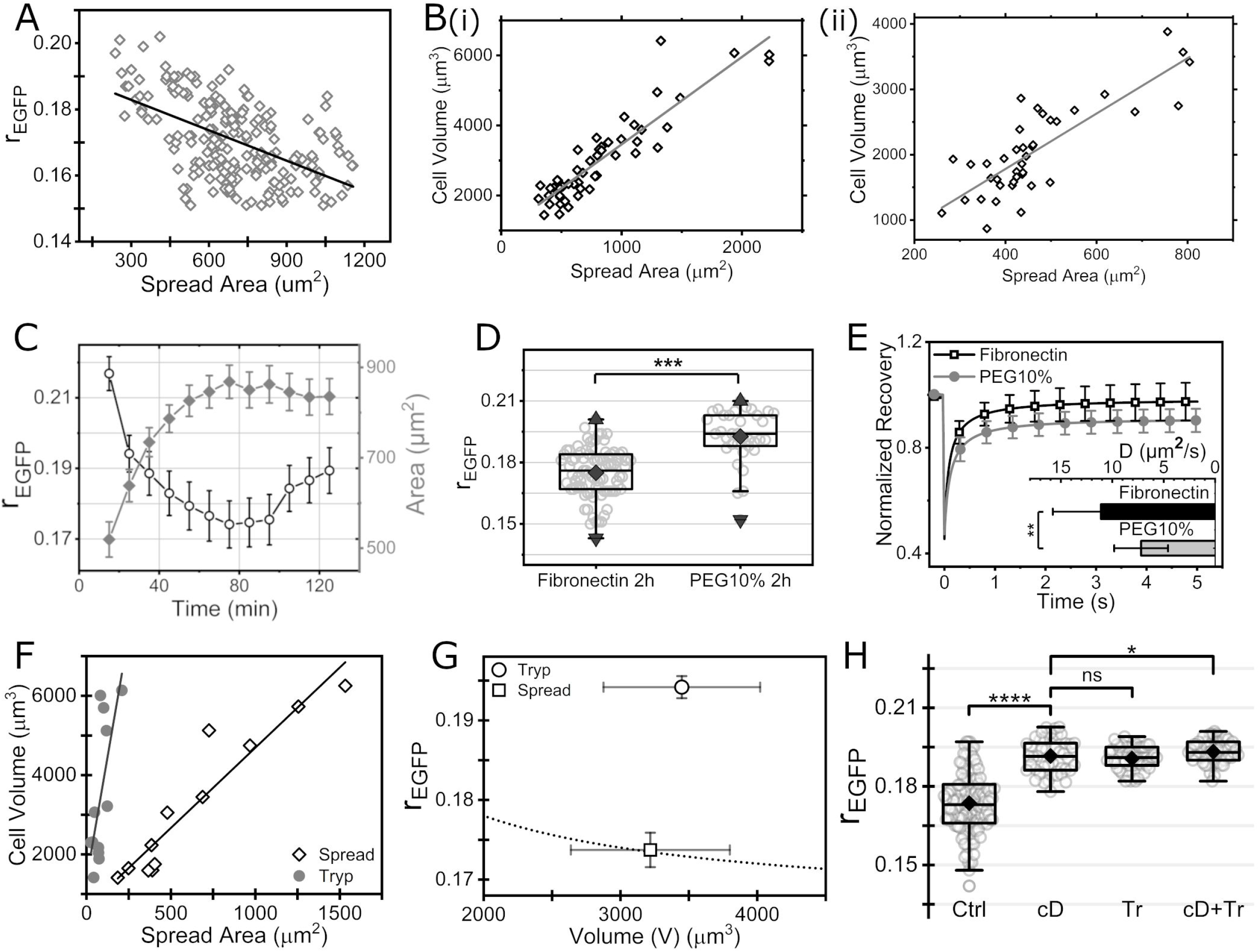
Cell spreading activates RVI. **(A)** Scatter plot and correlation of *r_EGF_* vs spread area of NIH/3T3-EGFP for cells seeded on Fibronectin. **(B)** Cell volume vs spread area of NIH/3T3-EGFP in well-spread condition **(i)** and during spreading **(ii)**. **(C)** *r_EGFP_* and spread area of NIH/3T3-EGFP while spreading on Fibronectin coated glass. **(D)***r_EGFP_* of NIH/3T3-EGFP seeded on Fibronectin or 10% PEG for 2 hours. **(E)** FRAP of NIH/3T3-EGFP seeded on Fibronectin or 10% PEG for 2 hours. **(F)** Cell volume vs spread area of NIH/3T3-EGFP before and after 30mins of trypsinization. **(G)** *r_EGFP_* vs cell volume (V) before and after 30mins of trypsinization **(H)** Comparison of *r_EGFP_* in NIH/3T3 control, Cytochalasin D (2μM,1h) treated, trypsinized, and trypsinized after Cytochalasin D treatment.

Interestingly, the area and volume in the population of spreading cells also exhibit a positive correlation (*r*=0.818). However, the *V* – *A* relation in spreading cells shows a slightly different slope: *V* = 2.5*A*+ 963.4 (**Fig 5B-ii**). It suggests that cell spreading dictates the spread area-MMC correlation in (**Fig 5A**) in NIH/3T3-EGFP.If cytoplasmic MMC were to be a sensor for cell volume regulation, cells must regulate their volume to maintain the MMC during cell spreading. On the contrary, we find that despite no change in the extracellular tonicity, the average cell volume increases, leading to a decrease in cytoplasmic MMC during cell spreading (**Fig 5C, Fig S5A**). There is a change in both cell volume and the cytoplasmic MMC in these time scales. This observation cannot establish whether the cells readjust their volume to regulate the cytosolic MMC or if the cell volume changes are indifferent to MMC changes. If the volume regulation is triggered to control MMC, then MMC must act as a sensor for volume regulation. To resolve this, we investigate the cytoplasmic MMC in cells seeded on surfaces that either promote (Fibronectin) or inhibit (10% PEG) cell spreading. Even after 2 hours of seeding, the cytoplasmic MMC in cells with compromised spreading is significantly higher than the well-spread cells (**Fig 5D, S5B**), establishing that cytoplasmic MMC does not trigger volume regulation in spreading cells; instead, the spreading state of the cells is necessary to initiate volume regulation. Cell spreading causes a significant decrease in the cytoplasmic MMC, causing a 5.4% change in cytoplasmic viscosity (η=4.87±1.15cP for cells on Fibronectin, η=7.5±2.06cP for cells on 10%PEG) (**Fig 5E**). In the reverse case of cell area contraction experiments, performed by exposing NIH/3T3-EGFP to Trypsin/EDTA, we observe insignificant change in the cell volume between the spread state or 30min after trypsinization. (**Fig 5F, S5C**), suggesting that contraction of cell spread area within few minutes, do not maintain the same *V* – *A* relation as in **Fig 5B**. Yet, we find measurable increase in cytoplasmic MMC during contraction, because of the disruption of cytoskeletal filaments to monomers, as we observe for actin (**Fig 5G, S5D**), and according to previous reports for microtubules (Celik *et al*., 2013). To test this, we first disassemble the actin filaments using Cytochalasin D and then initiate contraction by trypsin treatment in NIH/3T3-EGFP. Expectedly, the increase in *r_EGFP_* upon trypsinization is significantly less in Cytochalasin D treated cells when compared to control NIH/3T3-EGFP (**Fig 5H**).

### Plasma membrane tension acts as a sensor for cell volume regulatory apparatus

Cell volume regulatory apparatus can be broken down into three parts (i) cell volume sensor (ii) intracellular signaling cascades, which receives information related to change in volume and (iii) the volume regulatory action, leading to resetting of cell volume to a set-point. Regulatory volume change during osmotic imbalance involves the exchange of organic and inorganic osmolytes (Jentsch, 2016; Serra *et al*., 2021), which are smaller in size than the macromolecular crowders. Thus, there can be two possibly distinct features that could act as a volume sensor. 1) chemical signal, i.e., change in cytoplasmic MMC that alters the thermodynamic activity of the cytoplasm via excluded volume effect, and 2) mechanical signal, i.e., change in the mechanical properties of the membrane/cytoskeleton that alters the intracellular levels of osmolyte for enforcing regulatory volume change. (Dunham, 1995) proposed that the altered cytoplasmic MMC in hypertonic conditions acts as a sensor for volume regulation in RBC cells, suggesting that the changes in the thermodynamic activity of the cytoplasm are the sensor for volume regulation. These studies do not rule out the possibility that altered smaller-sized osmolyte level itself, which has a negligible effect on the thermodynamic activity, could be the sensor for cellular volume regulation during RVI. To identify whether cytoplasmic MMC acts as a volume sensor for the cell, we performed experiments that changed the cytoplasmic MMC without altering the levels of intracellular smaller-sized osmolytes. Hypertonic conditions involve the exchange of osmolytes (Roffay *et al*., 2021) and non-specific aggregation of proteins (**Fig 4**). As a result, the clustering of cytoplasmic proteins causes the *r_EGFP_* to rise above the expected curve (dotted line) (**Fig 6A**). Since Heclin treatment leads to an increase in *r_EGFP_*, whereas treatment of Cycloheximide or heat shock causes a decrement (**Fig 2H**), we investigated if the changes in *r_EGFP_* in these conditions is due to a change in crowder concentration or protein clustering. **Fig 4E** suggests no apparent clustering of cytoplasmic proteins in response to Heclin treatment. Thus, we conclude that Heclin treatment leads to a rise in cytoplasmic MMC. Next, we investigate whether the increase in cytoplasmic MMC in Heclin treated cells triggers RVI. **Fig 6B** establishes that there is no noticeable change in cell volume in response to increased (Heclin) or decreased (Cycloheximide) cytoplasmic MMC, demonstrating that changes in MMC in NIH fibroblast cells do not trigger RVI/RVD even until 4 hours of treatment. The cytoskeleton is the crucial component of the cell that decides its shape. We investigate its role in determining cell volume and cytoplasmic MMC. Disassembly of actin causes no noticeable change in cell volume; however, the cytoplasmic MMC increases (**Fig 6C, Fig 3D**). The rise in cytoplasmic MMC is not inversely proportional to the associated volume change because disruption of actin filaments leads to a significant increase in macromolecular crowder in the form of monomeric G-actin (**Fig 6C**, dotted line). This further indicates that change in cytoplasmic MMC caused by disassembly of actin filaments does not trigger RVI to achieve homeostasis of MMC. Interestingly, the disassembly of microtubules causes an increase in cell volume (**Fig 6C**). The associated change in MMC is inversely proportional to the volume change, indicating that Nocodazole treatment causes a change in cytoplasmic volume, and cells ignore the associated change in MMC. Cell volume changes simultaneously perturb plasma membrane organization (Roffay *et al*., 2021). To investigate the role of the plasma membrane behind cell volume regulation, we postulate the involvement of TNFR1 (Tumour Necrosis Factor Receptor 1), as it is reported to exhibit enhanced clustering and internalization during hypertonic stress (Rosette and Karin, 1996). TNFR1 is also known to localize in membrane caveolae (D’Alessio *et al*., 2010; Bae *et al*., 2019), and caveolae are the excess membrane folds responsible for mitigating membrane tension imbalances (Sinha *et al*., 2011; Roffay *et al*., 2021). Moreover, TNFR1 is upstream of NFκβ, which shows enhanced activity during hypertonic stress after 3-5h of stress induction (Roth *et al*., 2010; Farabaugh *et al*., 2020). Zafirlukast, a cysteinyl leukotriene antagonist (Zhou *et al*., 2019), is reported to inhibit TNFR1 clustering (Weinelt *et al*., 2021). Zafirlukast treatment abrogates RVI in NIH/3T3 in response to a hypertonic shock of 500mOsm (**Fig 6D**). In addition, Zafirlukast treatment leads to arrest of cell volume regulatory activity during spreading (**Fig 6E-F**) and microtubule depolymerized state (**Fig 6G**). To understand the effect of Zafirlukast treatment on the plasma membrane, we compare the membrane-fluctuation tension of control cells with that of Zafirlukast treated using interference reflection microscopy (IRM) (**Fig 6H**) (Biswas, Alex, and Sinha, 2017). Membrane fluctuation-tension closely follows membrane frame tension for an extensive range of tension values (Shiba, Noguchi, and Fournier, 2016). **Fig 6H-J** establishes that Zafirlukast treatment causes a rise in membrane tension. The amplitude of temporal fluctuations is decreased in cells treated with Zafirlukast (**Fig 6I**). Thus, increased membrane tension (**Fig 6J**) deprives the cells of their ability to trigger regulatory volume change in hypertonic, spreading, and microtubule depolymerized states.

**Fig 6:**
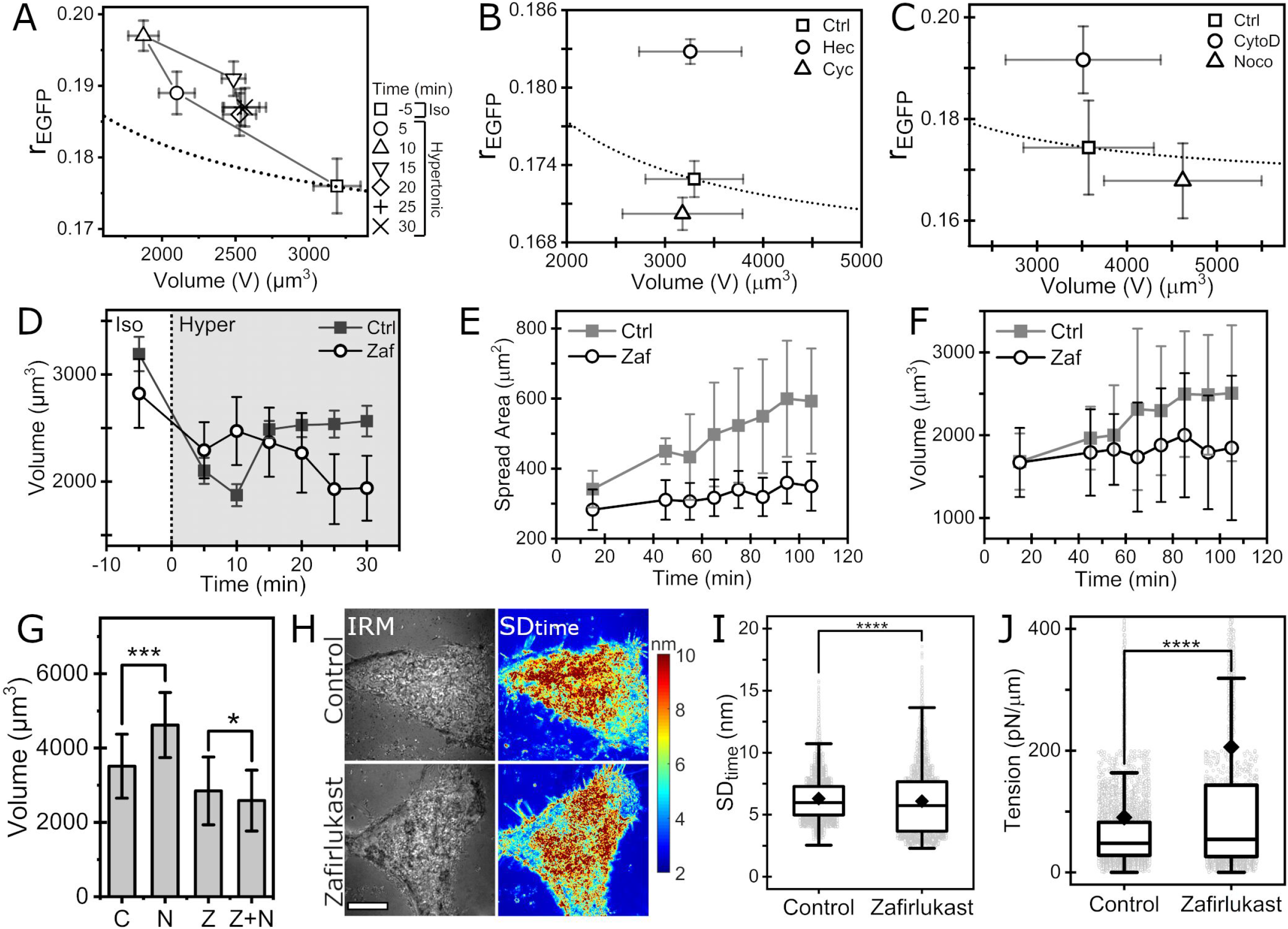
Plasma membrane tension acts as a sensor for cell volume regulatory apparatus. **(A)** Trajectory of *r_EGFP_* vs. cell volume at different time points after inducing hypertonic shock (500mOsm). Dotted line denotes the theoretical estimate of the trajectory. **(B)** *r_EGFP_* vs. cell volume after treatment with Heclin (70μM,4h) and Cycloheximide (1μM,4h) with respect to control in NIH/3T3-EGFP. **(C)** *r_EGFP_* vs. cell volume after treatment with Cytochalasin D (2μM,1h) and Nocodazole (20μM,1h) with respect to control in NIH/3T3-EGFP. **(D)** Cell volume trajectory in control and Zafirlukast (50μM,1h) treated NIH/3T3 in response to hypertonic shock (500mOsm, gray region). **(E-F)** Cell spread area trajectory **(E)** and corresponding cell volume trajectory **(F)** for control and Zafirlukast (50μM,1h) treated NIH/3T3 spreading on Fibronectin coated glass. **(G)** Comparison of cell volume in response to Nocodazole (20μM,1h) treatment in control (denoted C and N) and Zafirlukast (50μM,1h) treated (denoted Z and Z+N) NIH/3T3. **(H)** Representative Interference Reflection Microscopy images and corresponding *SD_time_* maps of vehicle control (DMSO) and Zafirlukast treated cells. Scale bar=10μm. **(I-J)** Comparison of *SD_time_* of membrane fluctuation **(I)** and tension **(J)** in vehicle control and Zafirlukast treated cells. (Control=6880 FBRs, 13 cells; Zafirlukast=4514 FBRs, 10 cells).

## Conclusion

Measurement of *r_EGFP_* in the cytoplasm and solutions of different BSA concentrations establish that *r_EGFP_* is a reliable indicator of macromolecular crowding in cells. MMC lowers *τ* and increases *r*_0_ of EGFP giving a linear relation between *r_EGFP_* and the concentration of the macromolecular crowder. We establish that cells of different lineages maintain spatially inhomogeneous and distinctly unique levels of average cytoplasmic MMC. The average cytoplasmic MMC in NIH-3T3 cells is maintained tightly for a relatively long time (at least 8h) if the spread area does not change significantly. Since cell-volume and cytoplasmic MMC are inversely related, measurement of *r_EGFP_* and cell volume allows us to explore the relation between cytoplasmic MMC and cell volume regulation. We find that heat-shock, trypsin-induced contraction, treatment of Cytochalasin D, Heclin, or Cycloheximide shifts the MMC in the cytoplasm without any change in cell volume. This observation establishes that the cells lack the ability for homeostasis of cytoplasmic MMC. Though the shift in MMC can alter a host of biochemical processes in the cytoplasm, NIH/3T3 cells lack the capabilities to regulate it. Imbalance in extracellular osmolarity is known to trigger RVI/RVD. Our manuscript identifies additional conditions not linked to osmolarity shift, such as cell-spreading and microtubule-depolymerized state of the cells in which cells invoke volume change autonomously. In these conditions, cells enforce volume changes, disregarding cytoplasmic MMC changes. Moreover, the volume and MMC are inversely related. Overall, our observations indicate that MMC does not act as a sensor for autonomous volume change by the cells. Next, we identified TNFR1 as a vital player required for invoking cell volume regulation. Inactivation of TNFR1 with Zafirlukast abrogates cellular ability for volume regulation. Having ruled out the role of cytoplasmic MMC in cell volume regulation, we studied the plasma membrane’s mechanical state (tension) in control and Zafirlukast treated cells. Zafirlukast treatment causes an increase in membrane tension and abolishes the ability of the cells to regulate their volume. Thus, this manuscript conclusively rules out the role of cytoplasmic MMC in sensing the cell volume. Instead, membrane tension plays a crucial role in activating cell volume regulatory apparatus in NIH/3T3 cells.

## Materials and methods

### Cell culture and pharmacological studies

NIH/3T3 cell line was procured from NCCS (National Center for Cell Science, Pune, India). RAW 264.7 cell line was a generous gift from Dr. Sanjay Dutta (CSIR-Indian Institute of Chemical Biology, Kolkata), while HeLa and MDA-MB-231 cell lines were kindly gifted by Dr. Prosenjit Sen (Indian Association for the Cultivation of Science, Kolkata). Fugene^®^ (Promega) was used to transfect cells with the following plasmids: pCAG-mGFP, a gift from Connie Cepko (Addgene plasmid # 14757); mCherry-Lifeact-7, a gift from Michael Davidson (Addgene plasmid # 54491); and EGFP-p65, a gift from Johannes A. Schmid (Addgene plasmid # 111190), following standard protocol. Cells cultured in DMEM (Himedia, #AI007G) at 37*°C,* 5% CO_2_ in a humidified incubator, were seeded on custom-made glass bottom 35mm petridishes. The glass was coated with 50mg/mL Fibronectin (Sigma, #F1141) to promote rapid adhesion and proper spreading, or with 10% PEG (Polyethylene glycol 1000, Sigma #81189) to arrest spreading in the appropriate cases. Before microscopy, cells were gently washed with 1xPBS twice, and culture media was replaced with phenol red-free DMEM (Gibco, #21063029), which could be supplemented with the required drug when necessary. For all pharmacological treatments, Cytochalasin D (Merck, #C8273), Nocodazole (Merck, #487928), Withaferin A (Merck, #W4394), Heclin (Tocris, #5433), Cycloheximide (Sigma, #18079), and Zafirlukast (Merck, #Z4152) were dissolved in DMSO, and working concentrations were reconstituted as indicated in appropriate places. For applying heat shock, cells were incubated at 42°C for 1h in the presence of 5% CO_2_. To create osmotic imbalance, cells were first imaged in isotonic complete medium, and then the culture media was replaced with either hypertonic or hypotonic complete medium using a custom-made flow system. Hypertonic media was prepared by adding Dextrose, Mannitol, or NaCl (Merck Empura) to phenol red-free DMEM at indicated concentrations and filtered for decontamination. Hypotonic media was prepared by adding autoclaved MiliQ water to phenol red-free DMEM at required amounts to create the desired osmolarity.

### *r_EGFP_* measurement

Cells seeded on glass-bottom petridishes were imaged with a 40x air immersion objective (NA 0.75) using the Zeiss AxioObserver Z1 epifluorescence microscope. Light from a mercury arc lamp (HXP 150) was passed through a linear polarizer (ThorLabs) to create horizontally polarized light. The resulting polarized fluorescence signal from the cells passes through a polarizing beam splitter (DV2, Photometrics) to divide the emission light into parallel and perpendicular polarizations. The light is then collected by a CMOS camera (Hamamatsu Orca Flash 4.0 C13440), and the polarized fluorescence signal appears as an image having 2048×2048 pixels, with each half (1024×2048 pixels) representing the parallel and perpendicularly polarized emission, respectively. Due to misalignment in the optical path, the two halves don’t completely overlap. To resolve this, fluorescent polystyrene microspheres of 200nm diameter are dried on a glass coverslip and imaged in the same arrangement as *r_EGFP_* measurement, such that the images of the beads may serve as fiduciary markers to register the two halves of the image. Using the Descriptor-based Registration plugin of Fiji (ImageJ) (Schindelin *et al*., 2012) and a custom Fiji Macro, the left (perpendicular channel, *I*_⊥_) and right (parallel channel, *I*_∥_) half of the 2048×2048 image is registered to create the best possible overlap of the corresponding pixels in both channels. Thence, *r_EGFP_* is calculated for each pixel using the relation:

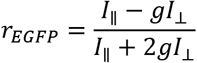

where *g* refers to the G-factor. Images of 100nM fluorescein solution are used to find the G-factor. To correct for background fluorescence, a 2048×2048pixel image of the phenol red-free DMEM, having no cells and illuminated by similar conditions as the experimental subjects, was subtracted from each 2048×2048 image. This process eliminates the background fluorescence of both the parallel and perpendicular channels in the correct ratio. The resultant *r_EGFP_* the image was saved as a 32-bit TIFF image file, thresholded based on intensity (15000-50000 count for 16-bit image), and further analyzed using a custom-written code in Fiji (ImageJ). Statistics of at least 30 cells were collected for each experiment, and their mean *r_EGFP_* and standard error of the mean were plotted. Confidence testing was performed with a Two-sample T-test in each case. Stars (*s) were assigned based on the general protocol for p-values.

### Cell volume measurement

Cells were imaged using the Zeiss LSM 780 light scanning confocal system using a 63x oil immersion objective. Z-stack images of 0.4μm step-size were taken using the AiryScan super-resolution mode to measure the whole cell volume. The whole Z-stack was binarized by an intensity threshold to mark the pixels containing cells as white and the background as black. Then, upon counting the number of white pixels contained in the binary Z-stack and multiplying the resultant with the appropriate voxel dimensions, the volume of the cells was calculated. Statistics of at least 10 cells were collected for plotting.

### *r_EGFP_* vs. *V* curve

The linear increment of *r_EGFP_* as a function of concentration (*C*) can be fitted to a straight line of slope *m* and Y-intercept *B*, which yields *C* = ^*r_EGFP_*^/*_m_* – *^B^/_m_*. Thus, the quantity *r_EGFP_* – *α*, where *α* = *^B^/_m_*, reflects a scaled measure of concentration. Following the relation *C*_1_*V*_1_ = *C*_2_*V*_2_, we get: (*r_EGFP_^—^ a*)*V* = (*r_EGFP,1-a_*)*V*_1_, and thus 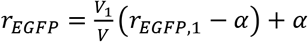.

### Fluorescence Recovery After Photobleaching (FRAP)

Using a home-built FRAP setup, photobleaching and recovery were imaged with a 488nm Laser (Coherent) through the 63x oil immersion objective of Zeiss AxioObserver Z1. Briefly, the 0.42mW Laser beam was split in a 90:10 ratio. The resultant beams were collimated using a lens system to be incident parallelly on the back focal plane of the microscope objective. The beams were aligned to illuminate the same spot (of 2μm diameter) when imaged with the 63x objective. The low-intensity beam was further dimmed using neutral density filters to minimize photobleaching and image the circular spot. To perform FRAP, the circular spot was continuously imaged at a rate of 50-100 frames per second with only the low-intensity beam. After 70-100 frames, the high-intensity beam was exposed for 10ms using a programmable shutter (Thorlabs, SC10) to achieve fast photobleaching. Imaging is continued for a total of 2000 frames, by which time the intensity of the spot becomes constant, indicating completion of recovery. The fluorophore’s diffusion rate and mobile fractions are calculated by fitting the intensity recovery data from the spot with a custom-written MATLAB code, as explained in (Kang *et al*., 2010). Before studying live cells, the FRAP setup was calibrated using a glycerol-water mix of known viscosity containing 100nM fluorescein (data not shown).

### Single Particle Tracking

Fluorescent polystyrene beads of diameter 200nm (Invitrogen, #F8888) were imaged with a 63x oil immersion objective at a rate of 100 frames per second to capture the thermal motion. For *in vitro* measurement, beads were suspended in BSA solutions at previously indicated concentrations. The beads were ballistically injected with the Helios Gene Gun (BioRad) delivery system for intracellular measurement. Cells were “shot” with a pressure of 100PSI from a distance of 3-4cm from the petridish. Cells were gently washed with serum-free media thrice to remove beads stuck on the plasma membrane or glass. Then the cells were incubated in phenol red-free DMEM at 37°C, 5%CO_2_ for 2h to allow them to recuperate. The trajectories of the fluorescent beads were extracted using the Mosaic plugin (Particle Tracking 2D/3D) of ImageJ. The following relation was used for MSD computation of a bead with trajectory (*x_t_,y_t_*): *MSD*(*τ*) = 〈(*x_t+τ_* - *x_t_*)^2^ + (*y_t+τ_* - *y_t_*)^2^〉, where, *τ* is the lag-time. MSD computation was performed using a custom-written MATLAB code.

### Calculation of viscosity (*η*) from Diffusion constant (*D*) or Mean Squared Displacement (*MSD*)

The Stoke-Einstein equation: *D* = *k_B_T*/6*πηr*, where *r* is the Stoke’s radius of the particle of interest, relates *D* with *η*. Using *r*=4.2nm for EGFP, we can calculate *η* for simple diffusion of EGFP. Since *MSD*(*τ*) = 4*Dτ* (in 2D), we can easily obtain *η* using the Stoke-Einstein relation.

### Time-resolved fluorescence anisotropy and fluorescence lifetime measurements

EGFP purified from *E. Coli* using Ni-NTA chromatography was lyophilized and reconstituted in Hepes (SRL, #63732) buffer of pH (7.2-7.6). The concentration of purified EGFP was estimated from UV absorbance and fluorescence. Subsequently, the reconstituted EGFP was diluted in appropriate solvents in a fixed ratio of 1:100 v/v for all experiments (except *r_EGFP_* vs EGFP concentration). Fluorescence lifetime and time-resolved anisotropy decay measurements were done using the DeltaFlex™ system (Horiba) using 4-side transparent UV quartz cuvettes.

### Fluorescence Lifetime Imaging Microscopy (FLIM)

FLIM was carried out using a pico-second 480nm laser (PicoQuant), and fluorescence lifetime data for individual pixels were fitted to mono-exponential decay using the SymPhoTime64 software. The resultant 32-bit TIFF image was analyzed in a similar way as in *r_EGFP_* measurements with Fiji (ImageJ) (Schindelin *et al*., 2012). Cells were seeded on glass-bottom 35mm petridishes, and hypertonic stress was applied following the same protocol as in *r_EGFP_* experiments. Data of 30 cells were used for statistics.

### Interference Reflection Microscopy

An inverted microscope (Nikon, Tokyo, Japan) with adjustable field and aperture diaphragms, 60x Plan Apo (NA 1.22, water immersion) with 1.5x external magnification, 100 W mercury arc lamp, (546 ± 12 nm) interference filter, 50:50 beam splitter, and CMOS (ORCA Flash 4.0 Hamamatsu, Japan) camera were used for IRM. Fast time-lapse images of cells were taken at 20 frames per second, and a total of 2048 frames were captured. Membrane fluctuations are quantified for regions within ~100 nm of the coverslip and termed First Branch Regions (FBRs). Calibration, identification of FBRs, and quantification of fluctuation amplitude (*SD_time_*)and tension were done as previously reported (Biswas, Alex and Sinha, 2017).

## Acknowledgements

We are grateful to the CSS and TRC confocal facility of the Indian Association for the Cultivation of Science for providing the confocal microscopy and TCSPC system. Special thanks to Dr. Aprotim Mazumdar and Mr. P. S. Kesavan of TIFR Hyderabad for assistance with the FLIM measurement, Mrs Debapriya Ghatak for assistance with confocal microscopy, Mr Subrata Das for assistance with TCSPC. We are also grateful to Dr Sanjay Dutta (CSIR-IICB, Kolkata), and Dr. Prosenjit Sen (IACS, Kolkata) for providing us with important cell lines.

## Author contributions and conflict of interest declaration

DKS, BS and PB designed the experiments and wrote the manuscript. PB prepared the figures; PB, DR, SJ and JD did the experiments. The authors declare no conflict of interest.

## Supplementary Figure Legends

**Fig S1:**
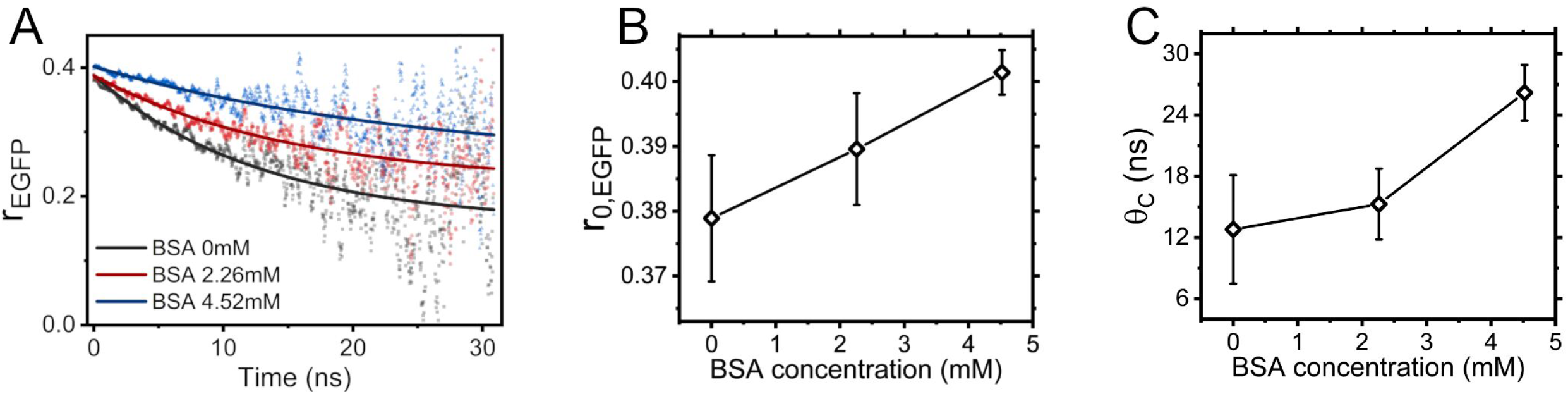
Intrinsic fluorescence anisotropy of EGFP (*r*_0_) increases with protein crowder concentration. **(A)** Time-resolved *r_EGFP_* in BSA solutions of 3 different concentrations prepared in Hepes buffer (pH 7.2). **(B)** The intrinsic fluorescence anisotropy of EGFP (*r*_0_) plotted as a function of crowder concentration. **(C)** Rotational correlation time (*θ*_C_) of EGFP plotted as a function of crowder concentration.

**Fig S2:**
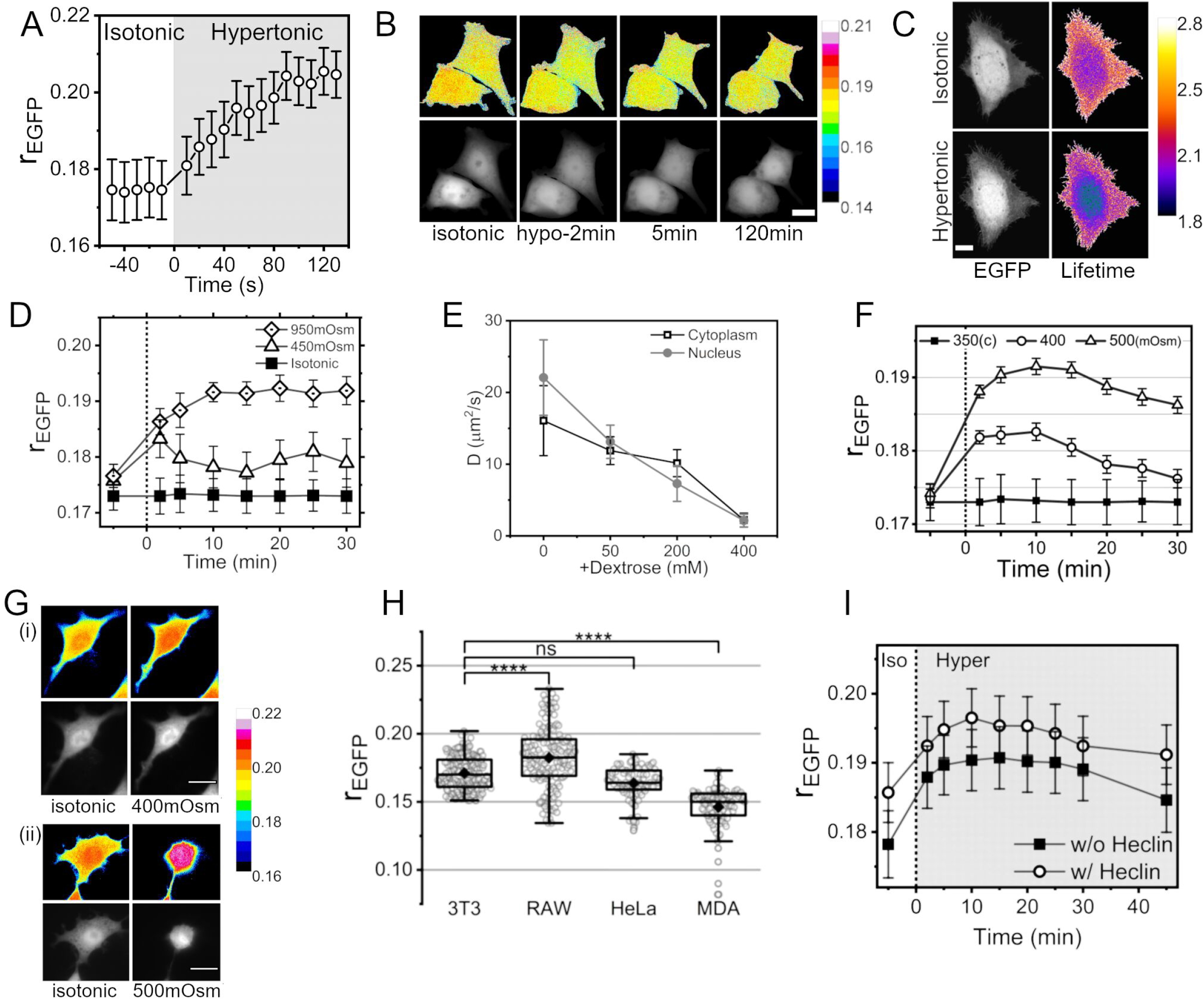
Cell’s ability to enforce RVI is cell lineage dependent. **(A)** Fast dynamics of *r_EGFP_* under 600mM hypertonic shock. **(B)** Time series of NIH/3T3-EGFP (*r_EGFP_* map-top row, intensity-bottom row) upon exposure to hypotonic media (170mOsm). Color bar shows the corresponding anisotropy values. Scale bar=15μm. **(C)** Fluorescence lifetime micrograph of cytoplasmic EGFP under hypertonic shock (grayscale-intensity image, Fire LUT-fluorescence lifetime image). Scale bar=15μm. **(D)** Cytoplasmic *r_EGFP_* upon hypertonic shock by NaCl. **(E)** Comparison of translational diffusion rate (D) of EGFP in the nucleus and cytoplasm in hypertonic conditions as measured by FRAP. **(F)** Hypertonic shock by mannitol yields a similar response as dextrose. **(G)** Intensity and *r_EGFP_* map of RAW264.7 cells exposed to 400mOsm **(i)** and 500mOsm **(ii)** hypertonicity. Scale bar=20μm. **(H)** Comparison of *r_EGFP_* of different cell lines under isotonic conditions. **(I)** Response of Heclin treated cells (open circles) to hypertonic shock (500mOsm) in comparison with untreated cells (filled squares).

**Fig S3:**
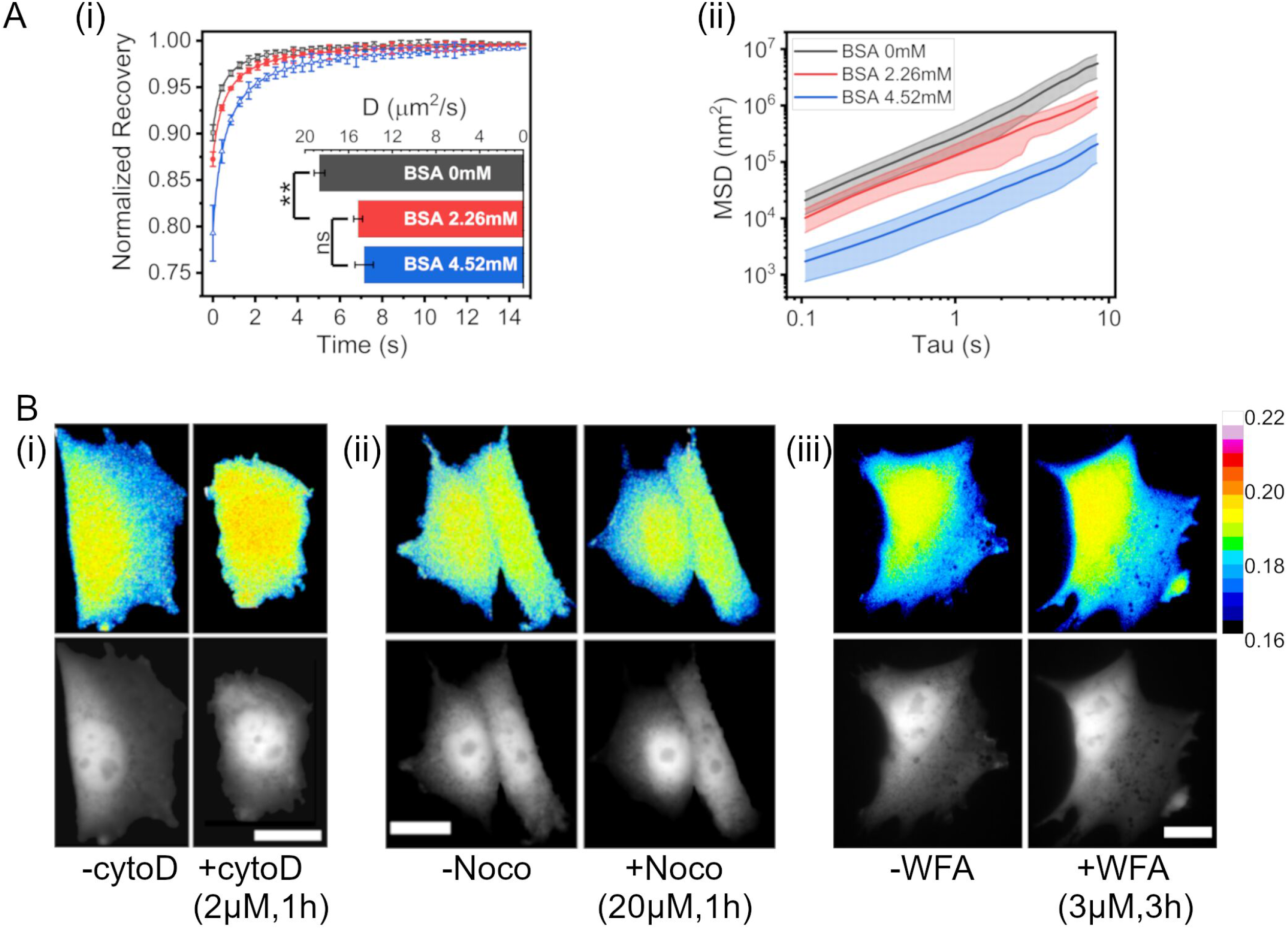
Despite diffusion, actin filaments hold spatially heterogeneous MMC in the cytoplasm. **(A)(i)** FRAP of EGFP done in BSA solutions of different concentrations. **(ii)** Single-particle tracking of fluorescent microspheres in the same BSA solutions. **(B)** Representative images (r_EGFP_-top row and intensity-bottom row) of NIH/3T3-EGFP after disruption of **(i)** actin by Cytochalasin D (2μM,1h), **(ii)** microtubules by Nocodazole (20μM,1h) and **(iii)** vimentin by Withaferin A (3μM,3h).

**Fig S4:**
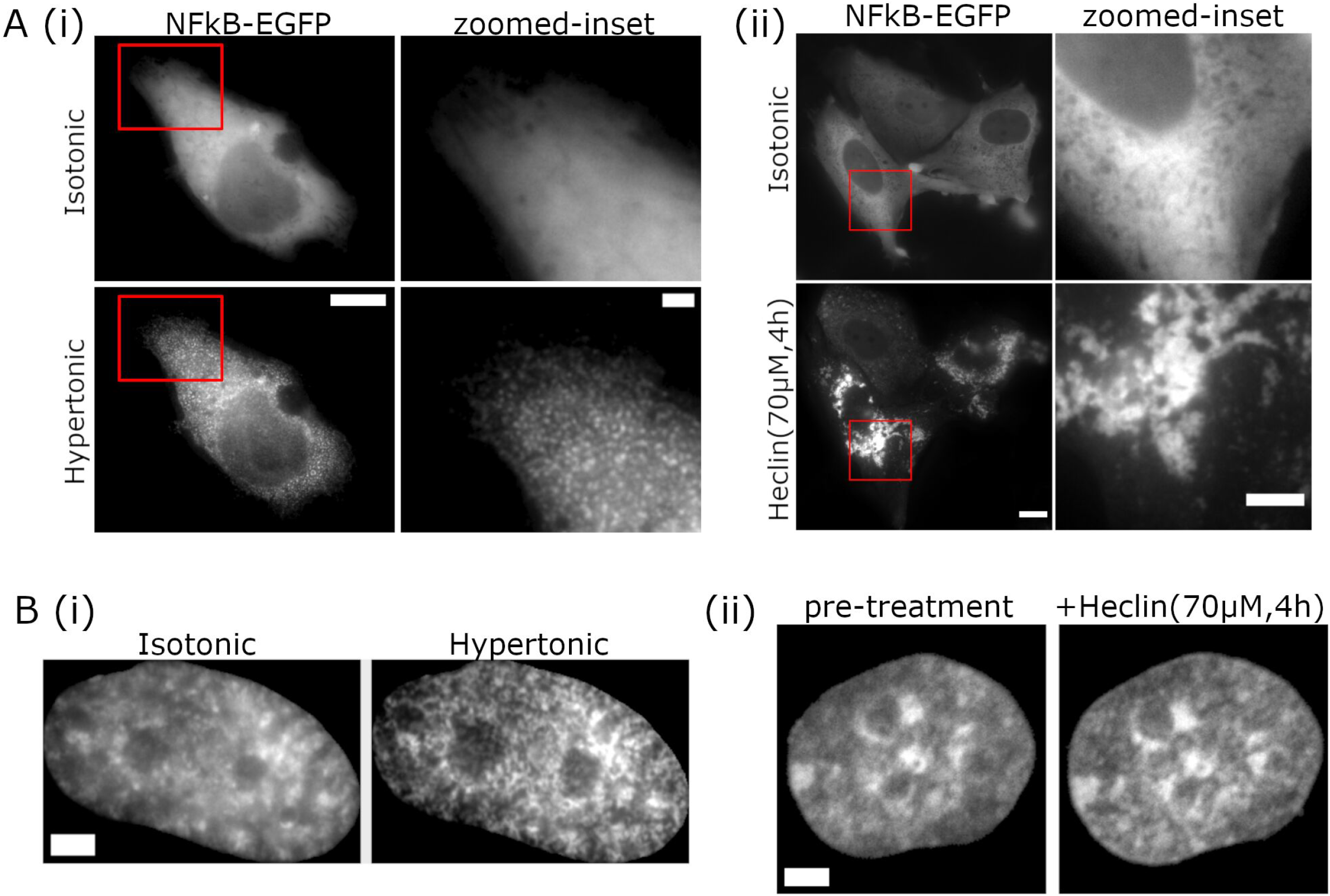
Hypertonic stress induces rapid and widespread submicron protein aggregation. **(A) (i)** NIH/3T3 show NFκβ-EGFP condensation upon hypertonic condition. Scale bar: main=10μm, zoomed inset=2μm. **(ii)** NFκβ-EGFP aggregates observed after treatment with Heclin (70μM,4h) in NIH/3T3. Scale bar=10μm, zoomed inset=5 μm (brightness-contrast of zoomed-inset was adjusted for better visualization) **(B)** DNA stained with Hoechst 33342 staining. **(i)** shows visible chromatin condensation during hypertonic (950mOsm) conditions. **(ii)** shows no apparent condensation of chromatin on Heclin treatment. Scale bar=5μm.

**Fig S5:**
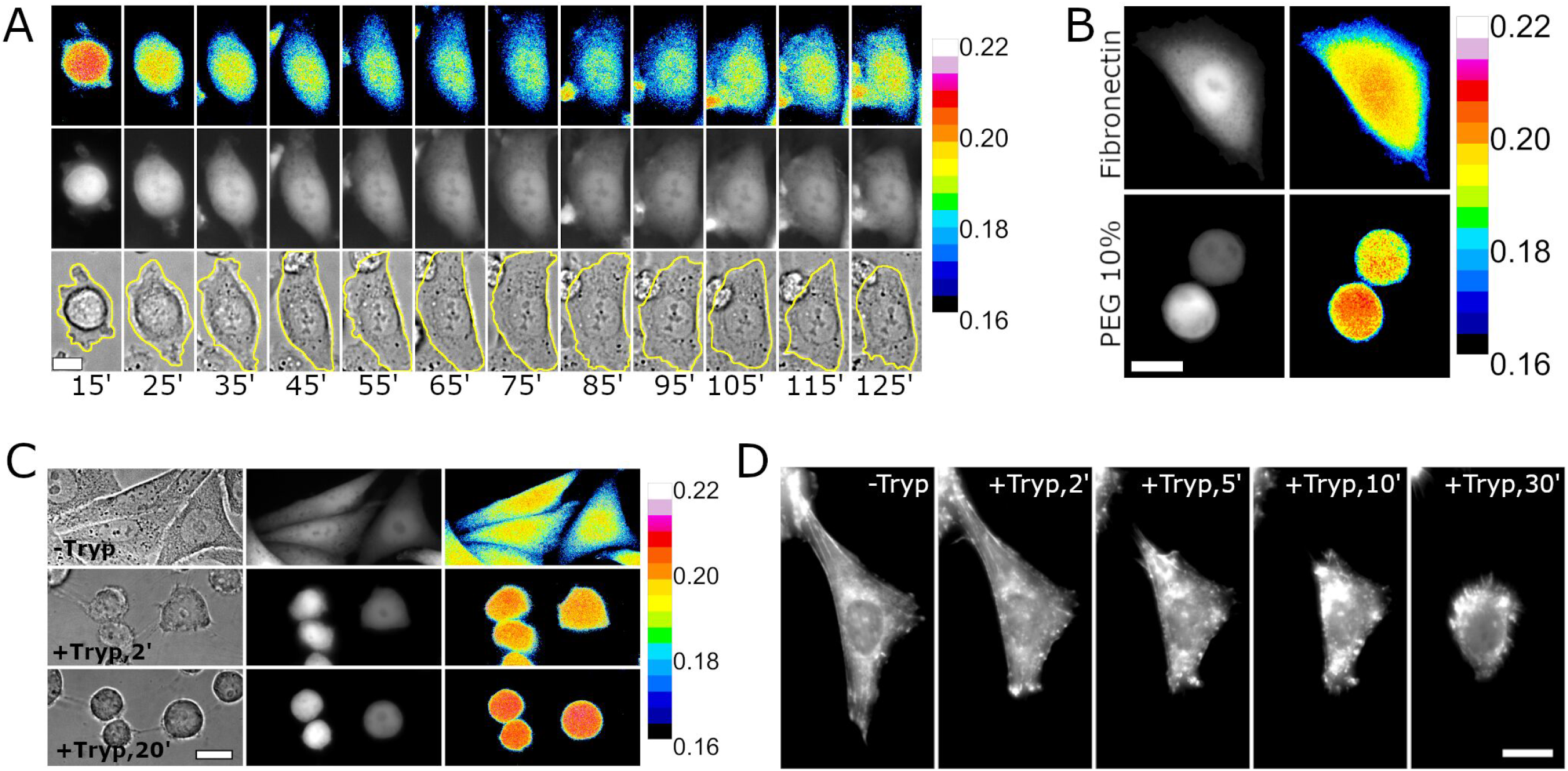
Cell spreading activates RVI. **(A)***r_EGFP_* map (top row), intensity (middle row), and bright-field (bottom row) of NIH/3T3-EGFP during spreading. **(B)** *r_EGFP_* and intensity of NIH/3T3-EGFP on Fibronectin and 10%PEG coated glass. **(C)** Bright-field, intensity and *r_EGFP_* map of NIH/3T3-EGFP undergoing deadhesion due to trypsinization. **(D)** Max intensity projection of LifeAct-mCherry expressing NIH/3T3 undergoing de-adhesion shows rapid disassembly of filamentous actin. Scale bar=10μm wherever shown.

